# ASTRID: Accurate Species TRees from Internode Distances

**DOI:** 10.1101/023036

**Authors:** Pranjal Vachaspati, Tandy Warnow

## Abstract

**Background:** Incomplete lineage sorting (ILS), modelled by the multi-species coalescent (MSC), is known to create discordance between gene trees and species trees, and lead to inaccurate species tree estimations unless appropriate methods are used to estimate the species tree. While many statistically consistent methods have been developed to estimate the species tree in the presence of ILS, only ASTRAL-2 and NJst have been shown to have good accuracy on large datasets. Yet, NJst is generally slower and less accurate than ASTRAL-2, and cannot run on some datasets.

**Results:** We have redesigned NJst to enable it to run on all datasets, and we have expanded its design space so that it can be used with different distance-based tree estimation methods. The resultant method, ASTRID, is statistically consistent under the MSC model, and has accuracy that is competitive with ASTRAL-2. Furthermore, ASTRID is much faster than ASTRAL-2, completing in minutes on some datasets for which ASTRAL-2 used hours.

**Conclusions:** ASTRID is a new coalescent-based method for species tree estimation that is competitive with the best current method in terms of accuracy, while being much faster. ASTRID is available in open source form on github.

## Background

Species tree estimation in the presence of gene tree incongruence is a major challenge for many biological analyses. Gene tree incongruence can result from a variety of processes, notably incomplete lineage sorting (ILS) [1], which is modelled by the multispecies coalescent (MSC) [2]. Concatenated maximum likelihood analyses is generally the most common method for species tree estimation from multiple loci, but can be statistically inconsistent, and even positively misleading, in some cases [3], thus converging to an incorrect tree with increasing amounts of sequence data. In recent years, a number of species tree estimation methods have been developed that are statistically consistent under the MSC, and so will converge in probability to the true species trees as the amount of data increases; see [4, 5, 6]. Methods that are statistically consistent under the MSC include ASTRAL [7], ASTRAL-2 [8], *BEAST [9], BEST [10], the population tree from BUCKy [11], METAL [12], MP-EST [13], NJst [14], SNAPP [15], STEAC [16], STEM [17], and SVDquartets [18]. While little is yet known about some of these methods (either because they have not yet been adequately studied or because they are not yet implemented), only a few of them (MP-EST, NJst, and ASTRAL-2) have been shown to be able to analyze very large datasets (especially those with large numbers of taxa) with high accuracy. MP-EST has been used more than either NJst or ASTRAL-2, but NJst is more accurate than MP-EST, and ASTRAL-2 is more accurate than both [8]. Furthermore, the currently available implementation of NJst is slower than ASTRAL-2, and cannot run on some datasets [8, 14].

In this paper, we present ASTRID, a new ILS-aware distance-based method for species tree estimation. Our approach is based on NJst, but is substantially faster, and, unlike NJst, functions even when each gene tree contains only a small portion of the data. The input to NJst is a set of unrooted gene trees. In the first step, an *n* × *n* matrix *D*[*x, y*] is computed, where *D*[*x, y*] is the average distance (in terms of number of edges) between *x* and *y* among all the gene trees. In the second step, neighbor joining [19], a very popular distance-based method of phylogeny estimation, is used to produce the species tree.

ASTRID improves on NJst by enabling other distance-based methods to be used in the second step. In particular, although NJ cannot be run on datasets with missing entries, other distance-based methods can, and ASTRID enables the use of these other methods. We also explore the use of more accurate distance-based methods. Thus, ASTRID is a very simple modification to NJst. As we will show, ASTRID is much faster than NJst.

The comparison between ASTRID and ASTRAL-2 and MP-EST, two established coalescent-based summary methods, is also interesting. ASTRID completed in minutes on some datasets where the other methods took hours, and was fast enough to analyze datasets with 1000 species and 1000 genes on a single processor within an hour (ASTRAL-2 and MP-EST take much more time on datasets of this size). Furthermore, ASTRID clearly dominates MP-EST in terms of accuracy, and is competitive with ASTRAL-2 (more accurate in some cases, and less accurate in others). Finally, ASTRID has desirable theoretical properties: it runs in polynomial time, and it remains statistically consistent under the MSC model without assuming the molecular clock, nor requiring rooted gene trees as input.

## Methods

### ASTRID

The input to ASTRID is a set of unrooted gene trees *T*_1_,…, *T*_*k*_. We let *S*_*i*_ = ℒ (*T*_*i*_) denote the leafset of *T*_*i*_, and *S* = ∪_*i*_ℒ (*T*_*i*_). Let |*S*| = *n*.

Step 1: Construct *n* × *n* matrix 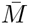:

1. For all *i* = 1, 2,…, *k*, compute *n* × *n* matrix *M*_*i*_, as follows. For pairs *p, q* of species where both are in *S*_*i*_, set *M*_*i*_(*p, q*) to be the number of edges in the path between *p* and *q* in *T*_*i*_. For all other pairs *p, q* (i.e., where one or both are not in *S*_*i*_), set *M*_*i*_(*p, q*) = 0. Thus, the only non-zero entries in *M*_*i*_ are for pairs of species in *T*_*i*_.
2. For all {*p, q*} ⊂ *S*, let *n*(*p, q*) be the number of trees *T*_*i*_ that contain both *p* and *q*.
3. Define *n* × *n* matrix 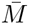 by setting 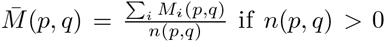 and 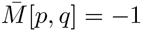 (to denote a missing value) otherwise.

Step 2: Compute tree on 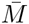 using a selected distance-based method

### Datasets

We tested species tree estimation methods on simulated datasets from previous publications, and also evaluated ASTRID on the mammalian biological dataset of 37 species, originally studied in [20]. Here we briefly describe the simulation procedures used to generate these datasets, and provide empirical statistics for the datasets in Table 1. See the original publications for details about the simulation protocols, and our supplementary online materials for links to the data.

**Table 1.** Empirical statistics of simulated datasets used in this study. The ILS level is measured by the average Robinson-Foulds distance (AD) between the true gene trees and the species tree, expressed as a percentage; ILS levels are then classified as low (L), moderate (M), high (H), or very high (VH).

| Dataset | # genes | # taxa | ILS level (AD%) | # sites | ref. |
| --- | --- | --- | --- | --- | --- |
| Avian very high ILS (0.5X) | 1000 | 48 | 60 (VH) | 500 | [22] |
| Avian high ILS (1X) | 1000 | 48 | 47 (H) | 250-1500 | [22] |
| Avian moderate (2X) | 1000 | 48 | 29 (M) | 500 | [22] |
| Mammalian high ILS (0.5X) | 200 | 37 | 50 (H) | 250-1000 | [22] |
| Mammalian moderate ILS (1X) | 200 | 37 | 29 (M) | 250-1000 | [22] |
| Mammalian low ILS (2X) | 200 | 37 | 21 (L) | 250-1000 | [22] |
| 10-taxon very high ILS | 200 | 10 | 89 (VH) | 100 | [23] |
| 10-taxon high ILS | 200 | 10 | 48 (H) | 100 | [23] |
| 15-taxon clocklike | 1000 | 15 | 82 (VH) | 100-1000 | [23] |
| ASTRAL-2 500K-1e6 (MC1) | 1000 | 200 | 69 (VH) | 300-1500 | [8] |
| ASTRAL-2 2M-1e6 (MC2) | 1000 | 200 | 33 (M) | 300-1500 | [8] |
| ASTRAL-2 10M-1e6 (MC3) | 1000 | 200 | 21 (L) | 300-1500 | [8] |
| ASTRAL-2 500K-1e7 (MC4) | 1000 | 200 | 68 (VH) | 300-1500 | [8] |
| ASTRAL-2 2M-1e7 (MC5) | 1000 | 200 | 34 (M) | 300-1500 | [8] |
| ASTRAL-2 10M-1e7 (MC6) | 1000 | 200 | 9 (L) | 300-1500 | [8] |
| ASTRAL-2 2M-1e6 (MC7) | 1000 | 10 | 17 (L) | 300-1500 | [8] |
| ASTRAL-2 2M-1e6 (MC8) | 1000 | 50 | 30 (M) | 300-1500 | [8] |
| ASTRAL-2 2M-1e6 (MC9) | 1000 | 100 | 34 (M) | 300-1500 | [8] |
| ASTRAL-2 2M-1e6 (MC10) | 1000 | 500 | 34 (M) | 300-1500 | [8] |
| ASTRAL-2 2M-1e6 (MC11) | 1000 | 1000 | 35 (M) | 300-1500 | [8] |

All datasets included both true and estimated gene trees, obtained by using maximum likelihood methods on the true sequence alignments, as well as species trees estimated on these gene trees obtained in the prior publications. Each gene tree had at most one copy of each species. We computed ASTRID species trees for these datasets, using various techniques for Step 2 (how to compute the species tree given the distance matrix).

We estimated the amount of ILS in the data by quantifying the average gene tree discord in the data, using the average Robinson-Foulds (RF) [21] distance between true gene trees and the model species tree, expressed as a percentage (written AD for “average distance”). We also explored some simulated datasets where the DNA sequence evolution was under the strict molecular clock. Model conditions with AD at most 25% can be considered low ILS, conditions with AD between 26% and 39% can be considered moderate ILS, conditions with AD between 40% and 59% can be considered high ILS, and conditions with AD of at least 60% can be considered very high ILS. In Table 1, we indicate these ILS levels for the different model conditions we study both with the AD value, but also the general level (L for low, M for moderate, H for high, and VH for very high).

#### Mammalian and avian simulated datasets

These datasets were created in [22] to evaluate method performance under model conditions similar to real data. Species trees were generated with MP-EST for the avian phylogenomics dataset with 48 species and 14,446 loci [24], and for a mammalian dataset with 37 species and 447 loci [25]. These species trees were used as basic model trees, with branch lengths in coalescent units. In addition, two other model species trees were created for each dataset by scaling the species tree branch lengths up (to reduce ILS) or down (to increase ILS). The ILS levels of the resultant model species trees were very heterogeneous, ranging from AD = 21% (low) to 50% (high) for the mammalian simulation, and from AD = 29% (moderate) to 60% (very high) for the avian simulation.

Both datasets had sequences of length 500 for all three model conditions. For the default (“1X”) branch length condition, the avian dataset also had sequences of length 250, 500, 100, and 1500, and the mammalian dataset had sequences of length 250, 500 and 1000. Sequence evolution on these datasets deviated from the strict molecular clock.

#### 10-taxon simulated datasets

These data were presented in [23], and explored two ILS levels (AD=48% (high) and AD=89% (very high)). Sequence evolution deviated from the strict molecular clock.

#### 15-taxon clocklike simulated datasets

These datasets evolved under a strict molecular clock, and were presented in [23]. The species tree was a caterpillar model tree (i.e., a path with leaves hanging off the path) with very short internal branches, and a long branch to the outgroup species. The ILS level in these data was very high (AD=82%).

#### ASTRAL-2 simulated datasets

These data were presented in [8], and provided a variety of model conditions with varying ILS levels, tree shapes, numbers of taxa, and sequence lengths per locus. SimPhy [26] was used to generate the species and gene trees, based on two parameters: the number of generations (given as the first number in the model) and the speciation rate (given as the second number). The number of generations simulated ranged between 500K, 2M, and 10M, and the speciation rate varied between 1e6 and 1e7. Model conditions with fewer generations had more ILS. Model conditions with the 1e6 speciation rate had speciation events nearer the tips (leaves) of the trees, while model conditions with the 1e7 speciation rate had speciation events nearer the root. The ILS levels varied from very low (AD = 9%) to very high (AD = 69%). Sequences evolved down the gene trees under multiple GTRGAMMA models that deviated from the strict molecular clock. Maximum likelihood gene trees were computed using FastTree-2.

#### Incomplete gene tree datasets

To explore performance on incomplete gene trees, we modified the ASTRAL-2 dataset by randomly removing taxa from trees in the 50-taxon datasets. Up to 40 taxa were removed from the 50-taxon dataset, and up to 5 taxa were removed from the 10-taxon dataset. In each of these cases, maximum likelihood gene trees were estimated using FastTree-2 version 2.1.7 SSE3 [27], using the following command:

~~~
fasttree -nt -gtr -quiet -nopr -gamma -n 1000 <fastafile> > <genetreefile>
~~~

 where 

~~~
<fastafile>
~~~

 was the input file of aligned sequences and 

~~~
<genetreefile>
~~~

 was the output file.

### Distance-based tree estimation methods

In order to explore the design space for ASTRID, we ran various distance-based methods for Step 2 (computing the tree from the distance matrix). For incomplete distance matrices (where some entries are -1, indicating that the pair of taxa do not appear together in any gene tree), we explored the methods in PhyD* [28]: *N J* *, *BION J* *, *M V R*\*, *U N J* *. These algorithms are all variants on neighbor joining that work on incomplete distance matrices. We also explored *F AST M E* [29], which is a heuristic for the minimum evolution problem.

### ASTRAL-2

To compute ASTRAL-2 species trees on the incomplete gene trees generated for the ASTRAL-2 datasets, we ran ASTRAL-2 version 4.7.8, using command line arguments

~~~
java -Xmx4000M -jar astral.4.7.8.jar -i <genetrees> -o <outputtree>
~~~

### Computing tree error

All trees computed in this study were fully resolved. We report the RF tree error (the proportion of the branches in the model tree missing from the estimated tree), using scripts that are available in the supplementary online materials.

## Results

### Selection of distance-based tree estimation method for Step 2

First, we evaluated various distance-based tree estimation methods to determine which one would be most accurate for the tree computation phase of ASTRID. Results on datasets with all complete gene trees (no missing species in any gene) are shown in Figure 1 and results on datasets with incomplete gene trees are shown in Figure 2. Note that for datasets with entirely complete gene trees, FastME performed as well as or better than the other distance-based methods, but there were datasets with incomplete distance matrices in which FastME had very poor accuracy. Therefore, we selected FastME to analyze datasets where the distance matrix has no missing entries, since it had the best accuracy. For the datasets with incomplete distance matrices 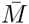 (indicated by 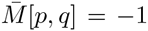 for some p,q), we selected BioNJ*, since it generally had among the most accurate results of these PhyD* methods.

**Figure 1.**
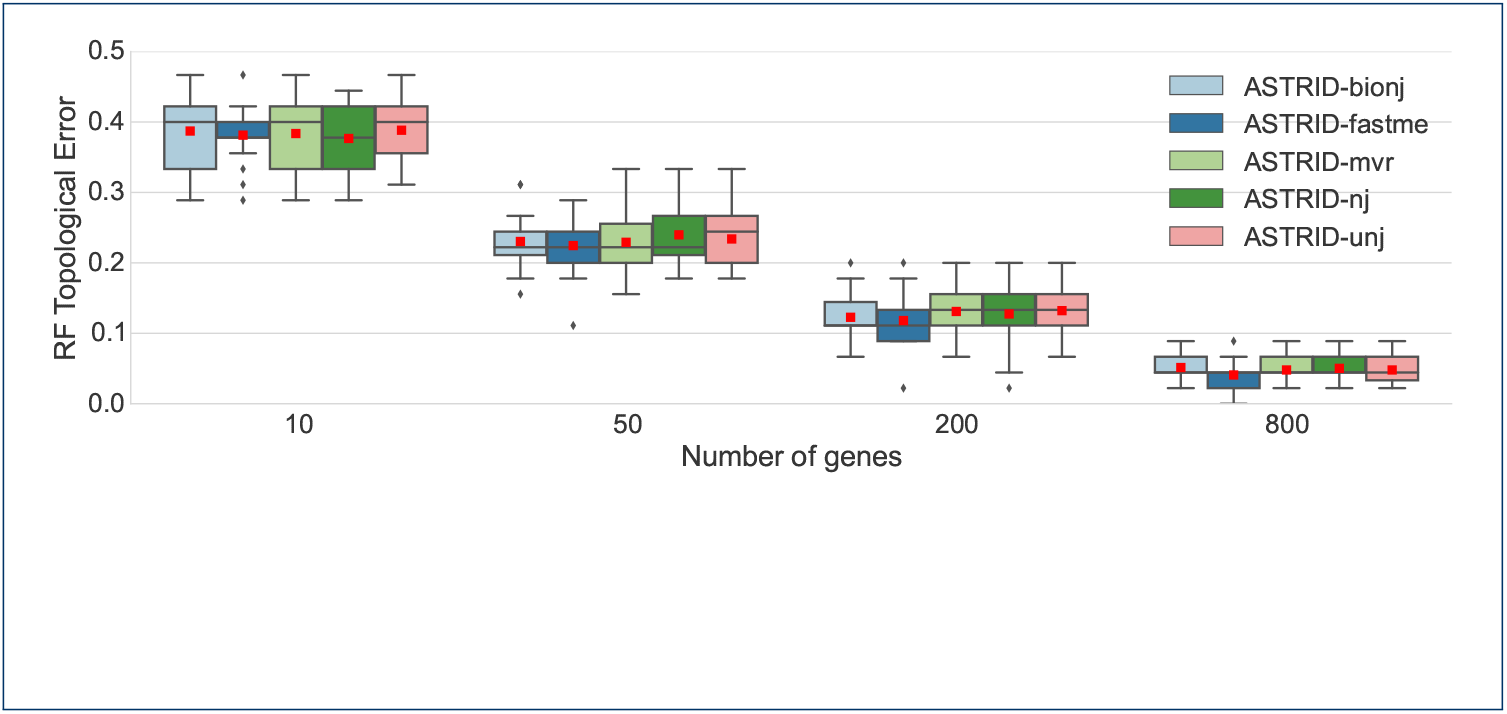
A comparison of ASTRID variants on the moderate ILS avian simulated datasets with 500bp, using different distance-based methods for the tree estimation phase. We report RF topological error rates over 20 replicates. Red dots represent means, while lines represent medians and boxes represent quartiles.

**Figure 2.**
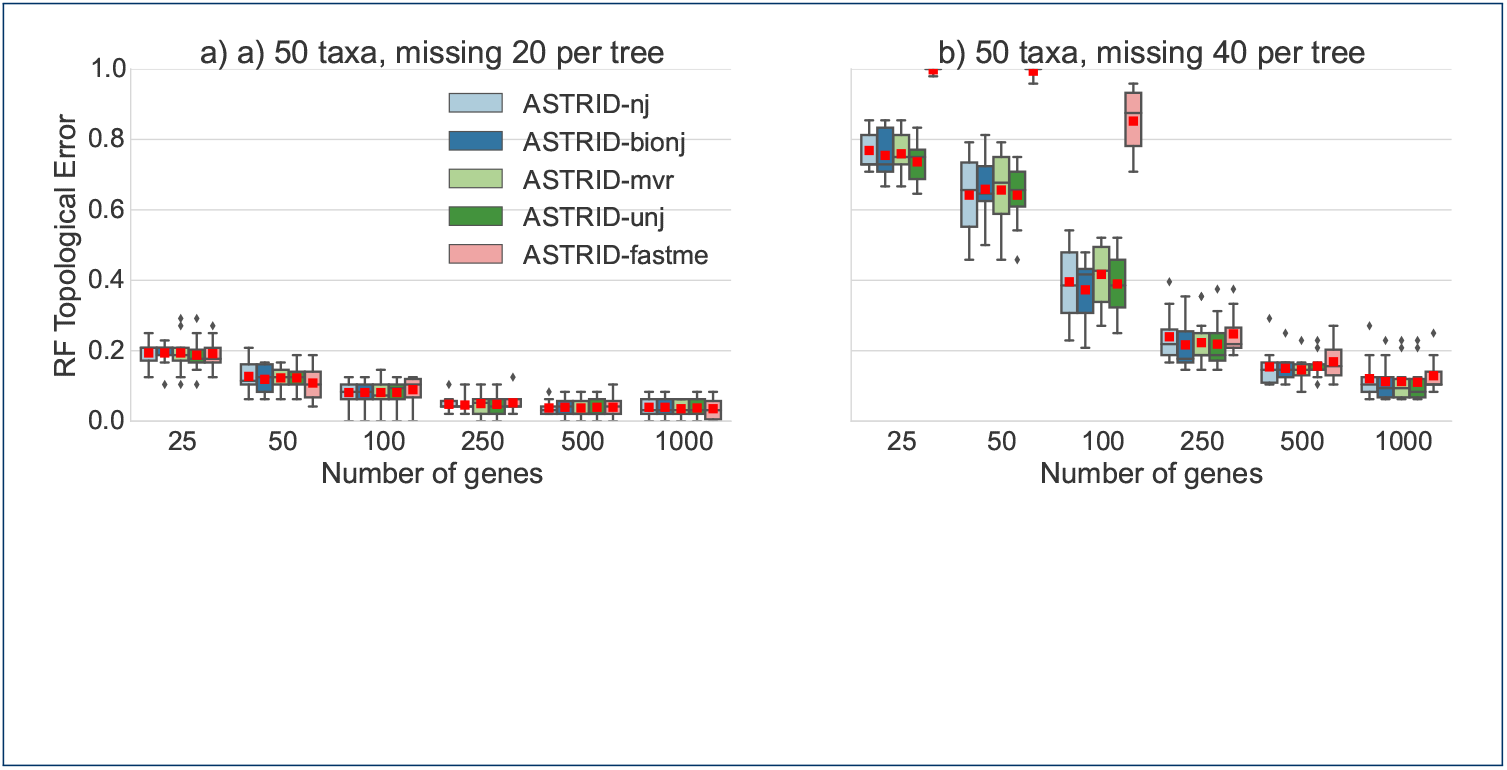
Comparison of ASTRID variants on 50-taxon ASTRAL-2 MC8 datasets with missing taxa. We show average RF error rates over 50 replicates for ASTRID variants, that differ in terms of the method used to compute the tree from the distance matrix. The datasets have taxa randomly removed from each gene and the sequence lengths truncated to 300bp. Red dots represent means, while lines represent medians and boxes represent quartiles.

### Comparison of ASTRID, ASTRAL, and MP-EST

We begin with a comparison between ASTRID, ASTRAL-2, and MP-EST on the avian simulated datasets with high (1X) ILS, varying number of genes and sequence alignment lengths, but where all genes are complete; see Figure 3.

**Figure 3.**
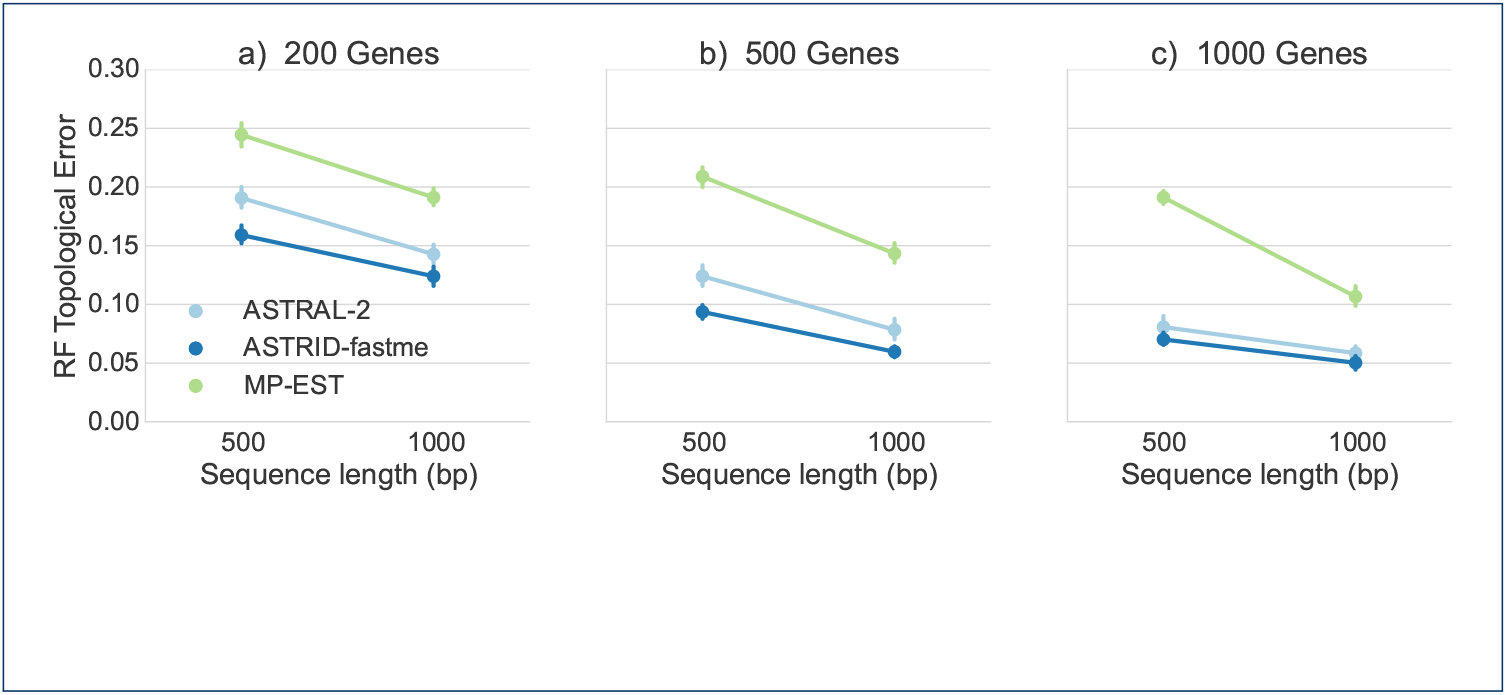
Comparison of ASTRID, ASTRAL-2, and MP-EST on the avian simulated data. The simulated data evolve under 1X (high ILS) species tree branch lengths, and with varying gene sequence lengths. We report mean RF rates with standard error bars over 20 replicates.

All methods improved with increasing numbers of genes or increasing sequence length; however, the methods differed substantially in terms of their accuracy. Across all conditions we explored, MP-EST had the highest error and ASTRID had the lowest error. ASTRAL-2 was in between, but was closer to ASTRID than to MP-EST. The gap between MP-EST and ASTRID was very large, and increased with the number of genes. For example, at 1000 genes and gene sequence alignments of length 500, MP-EST had 19% RF error while ASTRID had about 7% RF error. The gap between ASTRID and ASTRAL-2 was substantial on the 200- and 500-gene cases, but very small on the 1000-gene case.

Thus, although MP-EST is statistically consistent under the MSC model and hence theoretically robust to ILS, it did not have particularly good accuracy on these data. Among all coalescent-based methods, MP-EST is probably the one that has been used the most in biological data analyses, but its performance here and in [8, 30] demonstrates that it is not competitive with the best methods on datasets with even moderate numbers of species. Therefore, we omit MP-EST from the rest of this study.

### Comparison of ASTRID and ASTRAL-2 on complete gene trees

#### Comparison on avian datasets

Figure 4 shows the performance of ASTRAL-2 and ASTRID on avian simulated datasets under three ILS conditions (moderate, high, and very high). Both methods performed better when provided with more genes, and both performed worse on higher levels of ILS. Overall, ASTRID tended to outperform ASTRAL-2, with the largest effect seen when many genes were available. With 800 genes available, the ASTRID species tree had a RF error rate that was 2.4 percentage points better than ASTRAL-2’s under the very high and high ILS model conditions, and 1.2 percentage points better for the moderate ILS model condition. On the moderate ILS model condition, ASTRID had the greatest advantage over ASTRAL-2 for moderate numbers of genes. Above 200 genes, the error rate dropped below ten percent for both ASTRAL-2 and ASTRID, and ASTRID had an average advantage of only about one percentage point.

**Figure 4.**
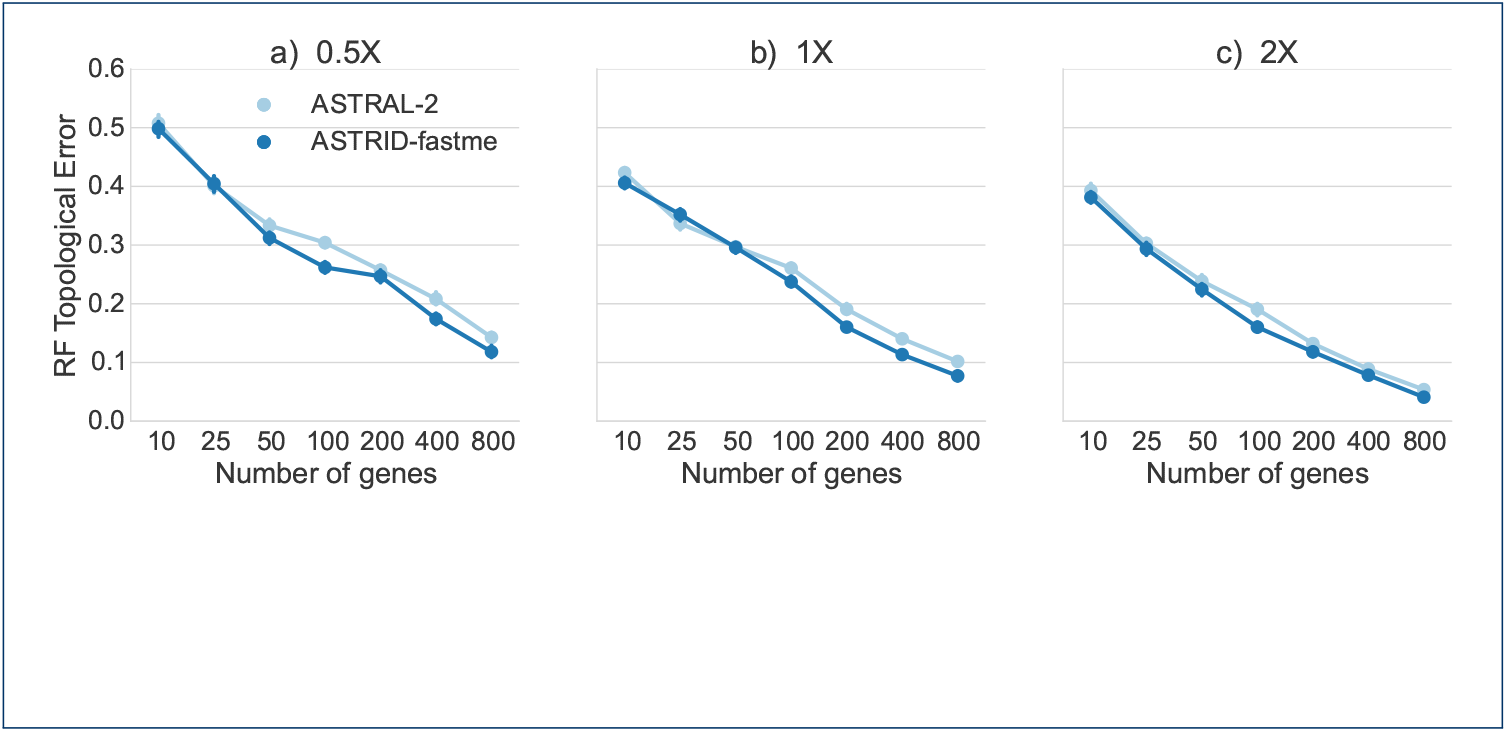
Comparison of ASTRID and ASTRAL-2 on avian simulated datasets. We show average RF error rates and standard error bars for 20 replicates. Gene sequence alignments have 500 sites and varying amount of ILS. Species tree branch lengths of 0.5x have very high ILS, 1X has high ILS, and 2x have moderate ILS.

It is well known that summary methods improve in accuracy as the number of sites per gene or the number of genes increase [31, 32, 33, 34]. We explored the impact of varying the sequence length and number of genes on the avian datasets with high (1X) ILS, as well as on true gene trees. Figure 5 shows results on 10, 100, and 1000 genes; results on other numbers of genes have the same trends (data provided in supplementary materials). As expected, both methods improved with increased sequence length, and had their best accuracy on true gene trees. Both methods also improved as the number of genes increased. ASTRID was always at least as accurate as ASTRAL-2, with the biggest improvement for shortest sequences (with 250bp).

**Figure 5.**
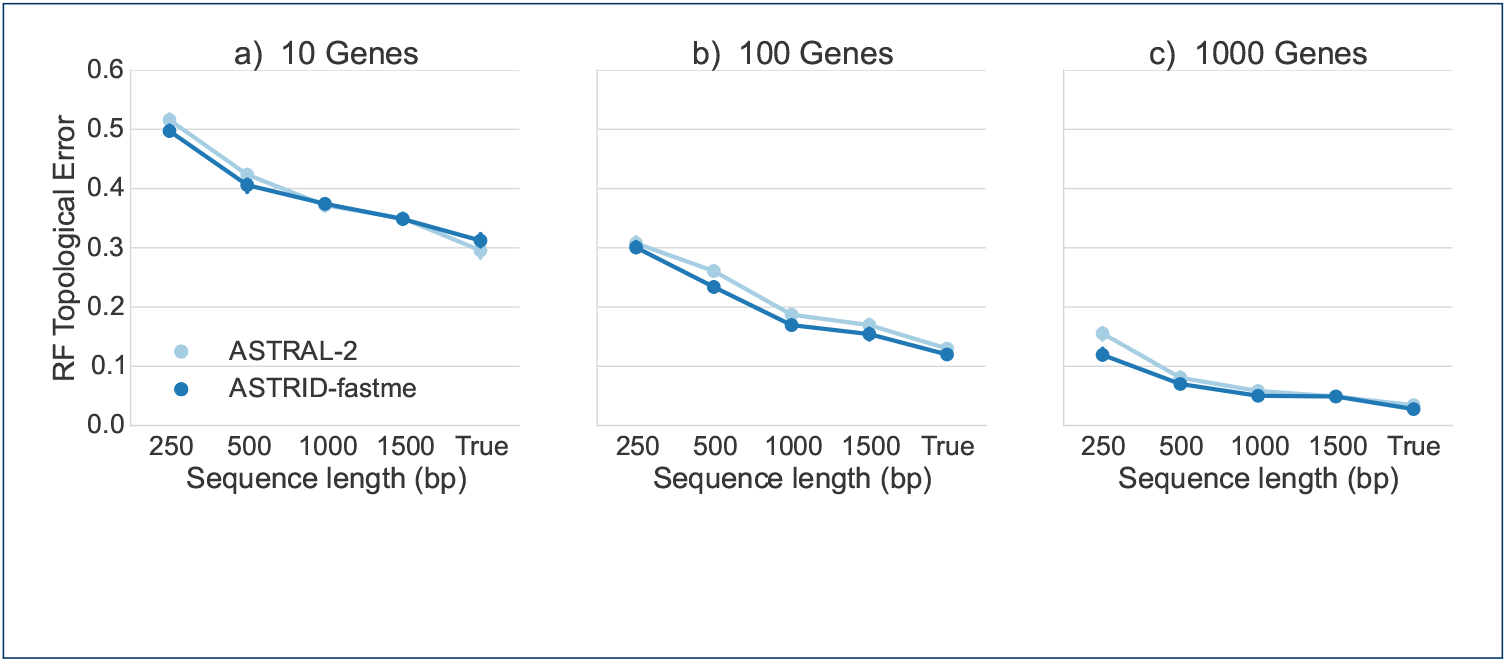
Performance on the avian simulated data with 1X species tree branch lengths, varying gene sequence length and number of genes. We report RF rates over 20 replicates.

#### Comparison on mammalian datasets

A comparison of ASTRAL-2 and ASTRID on the mammalian datasets with different levels of ILS (high, moderate, and low) is given in Figure 6. ASTRAL-2 and ASTRID performed fairly similarly on the low (2X branch lengths) and moderate (1X branch lengths) ILS conditions. Under the high ILS level (0.5X branch lengths), ASTRAL-2 was fairly consistently more accurate than ASTRID, with the largest improvement on the 10-gene case.

**Figure 6.**
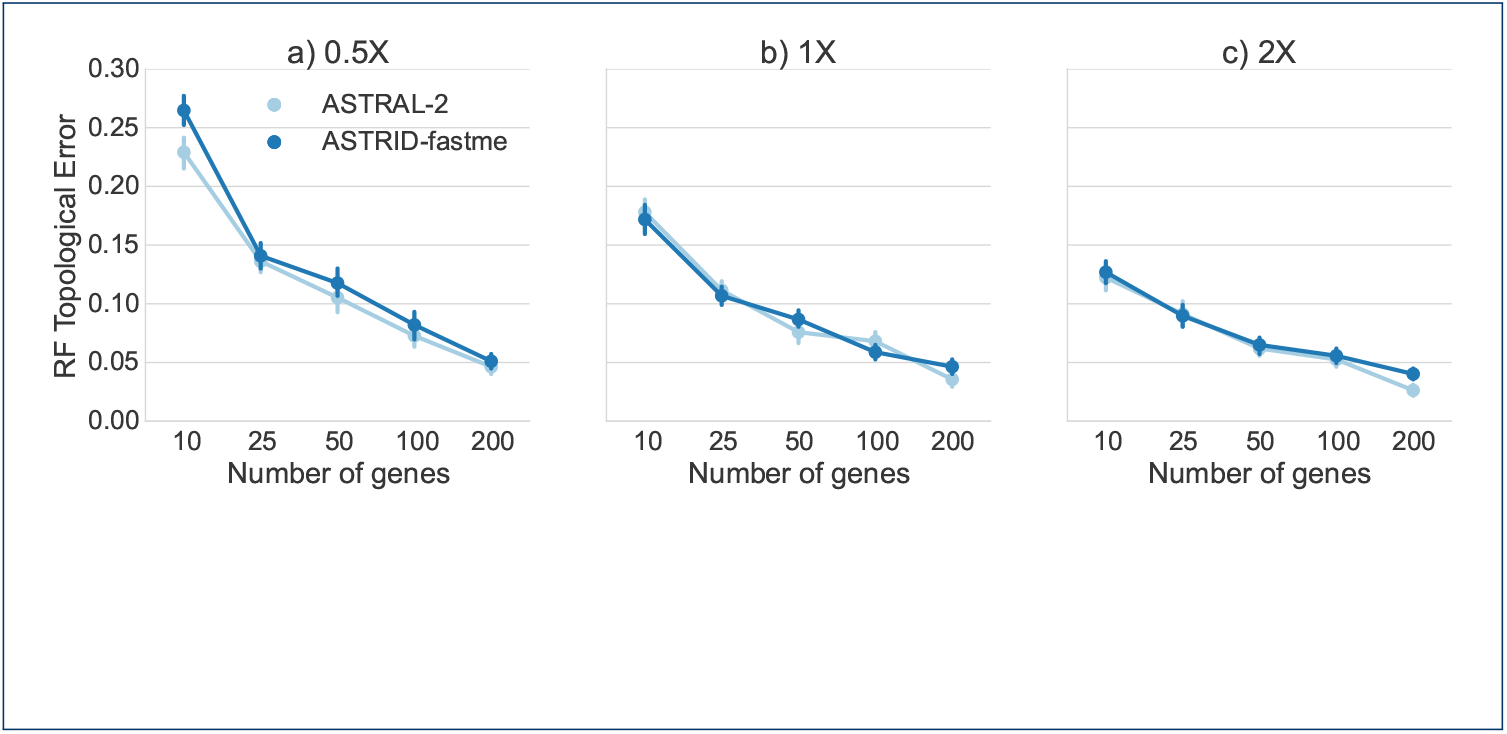
Comparison of methods on mammalian simulated datasets, varying ILS level and number of genes. We show average RF error rates and standard error bars for 20 replicates. Gene sequence alignments had 500 sites. Model conditions varied in ILS level from high (0.5x branch lengths) to low (2X branch lengths).

#### Comparison on the ASTRAL-2 datasets

We explored performance on the ASTRAL-2 datasets with 200 taxa (model conditions MC1 to MC6, see Fig. 7). These model trees varied in ILS level, with MC1 and MC4 having very high ILS, MC2 and MC5 having moderate ILS, and MC3 and MC6 having low ILS. Under MC2, MC3, and MC5, the two methods had essentially identical accuracy. However, under MC1, MC4, and MC6, ASTRAL-2 had an advantage over ASTRID. In MC1 and MC4, the improvement disappeared at 100 genes, but in MC6 ASTRAL-2 was still more accurate than ASTRID on 100 genes.

**Figure 7.**
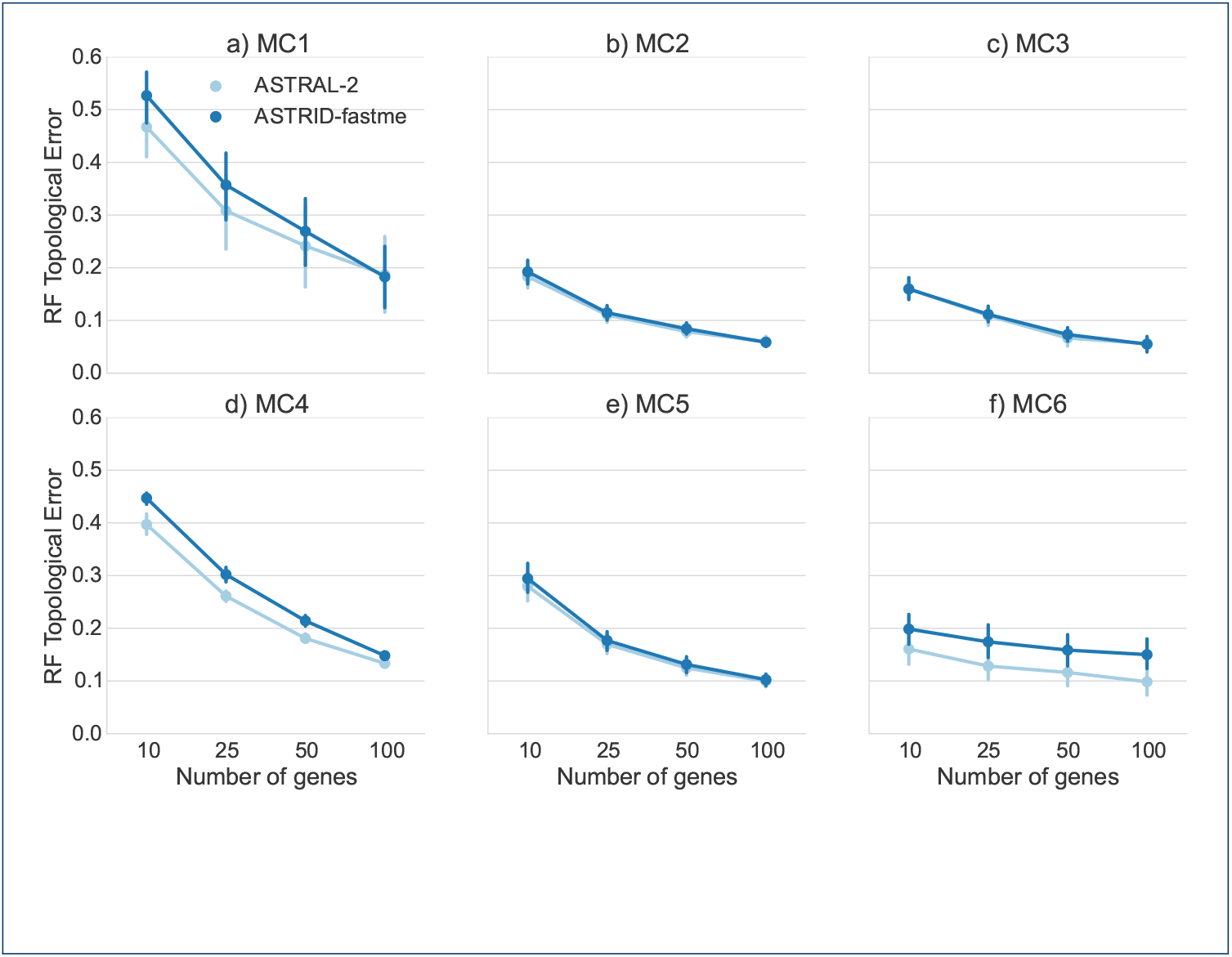
Comparison of ASTRID and ASTRAL-2 on the simulated ASTRAL-2 datasets with 200 taxa, varying levels of ILS, tree shape, and number of genes. We report RF error rates and standard error bars over 10 replicates. See Table 1 for information on the model conditions listed.

#### Comparison on the 15-taxon datasets

The 15-taxon datasets evolved on a caterpillar species tree under very high ILS (AD=82%), the highest ILS considered in this study. We explored performance under two sequence lengths (100bp and 1000bp) and varied the number of genes from 10 to 1000. Results on the 15-taxon datasets (Fig. 8) showed very close performance between ASTRID and ASTRAL-2. On the 100bp alignments and on 1000bp alignments with at least 100 genes, the two methods could not be distinguished. However, on 1000bp alignments with at most 50 genes, ASTRAL-2 had an advantage over ASTRID.

**Figure 8.**
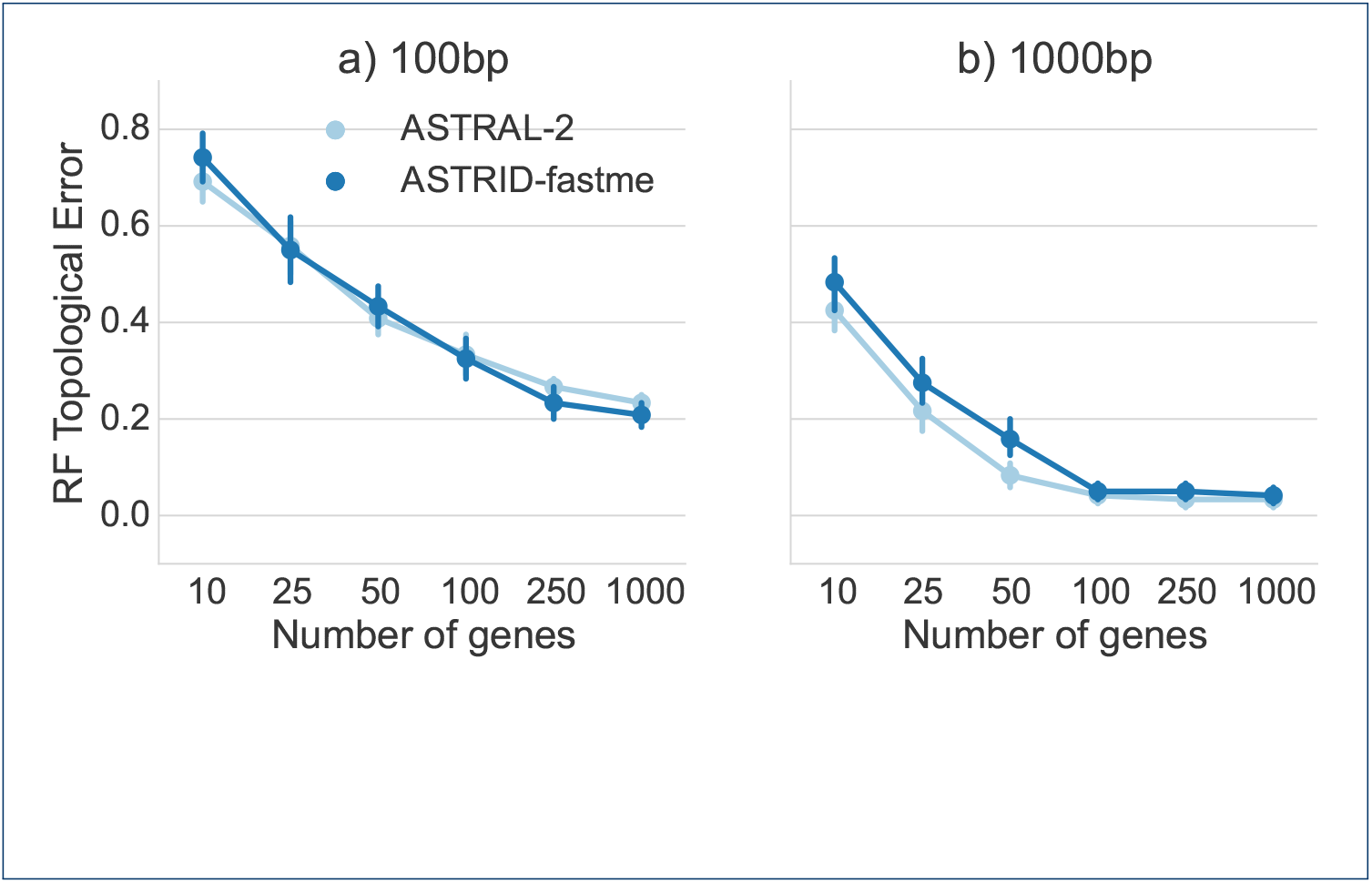
A comparison of ASTRID and ASTRAL-2 on the 15-taxon simulated datasets for two different sequence lengths. The 15-taxon datasets evolve down gene trees generated by a caterpillar tree with very high ILS (AD=82%), the highest ILS condition explored in this study.

#### Comparison on the 10-taxon datasets

The 10-taxon datasets evolved under two different ILS levels - high and very high, and we explored performance on both true and estimated gene trees; see Figure 9. In general, ASTRID and ASTRAL-2 had very close accuracy on these data, but there were some cases where they had different accuracy levels. For example, on the high ILS condition with estimated gene trees, ASTRAL-2 was more accurate than ASTRID for 200 genes, and ASTRID was more accurate than ASTRAL-2 on 25 genes.

**Figure 9.**
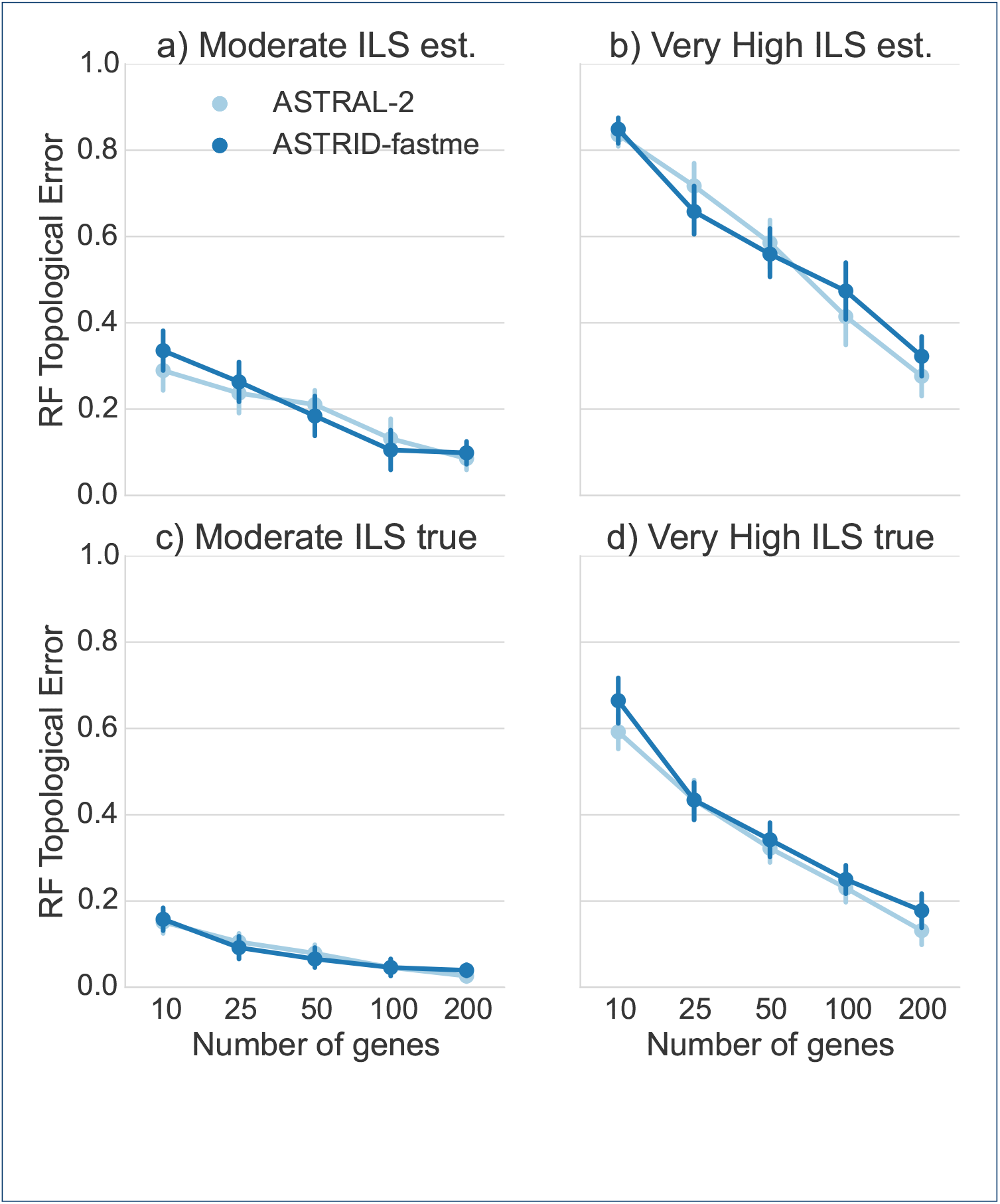
Results on true and estimated gene trees on 10-taxon datasets with two ILS levels (high and very high) All gene sequence alignments have 100bp. We report RF rates and standard error bars over 20 replicates.

### Performance on incomplete gene trees

We explored the impact of missing data on ASTRAL-2 and ASTRID by deleting taxa from gene trees in the 50-taxon datasets (MC8) from the ASTRAL-2 collection, using 150bp per gene, and varying the number of genes and the amount of missing taxa; see Figure 10. ASTRAL-2 and ASTRID had very similar topological accuracy throughout these experiments. With low amounts of missing data (20% to 40% missing taxa from each gene tree), both methods had very good accuracy (below 5% tree error) by 500 genes. With 60% of the taxa missing from each gene tree, the error rates increased for low numbers of genes (above 20% RF error for up to 100 genes), but then declined to about 10% by 1000 genes. With 80% of the taxa missing from each gene (so that all gene trees have only 10 taxa out of 50), error rates were very high with 25 genes (at least 85% RF), but decreased quickly with increases in the number of genes, so that at 500 genes the error rate was 24%, and then at most 18% at 1000 genes. The trends suggest that the error rates had not plateaued, and that adding additional incomplete gene trees should result in continued improvement.

**Figure 10.**
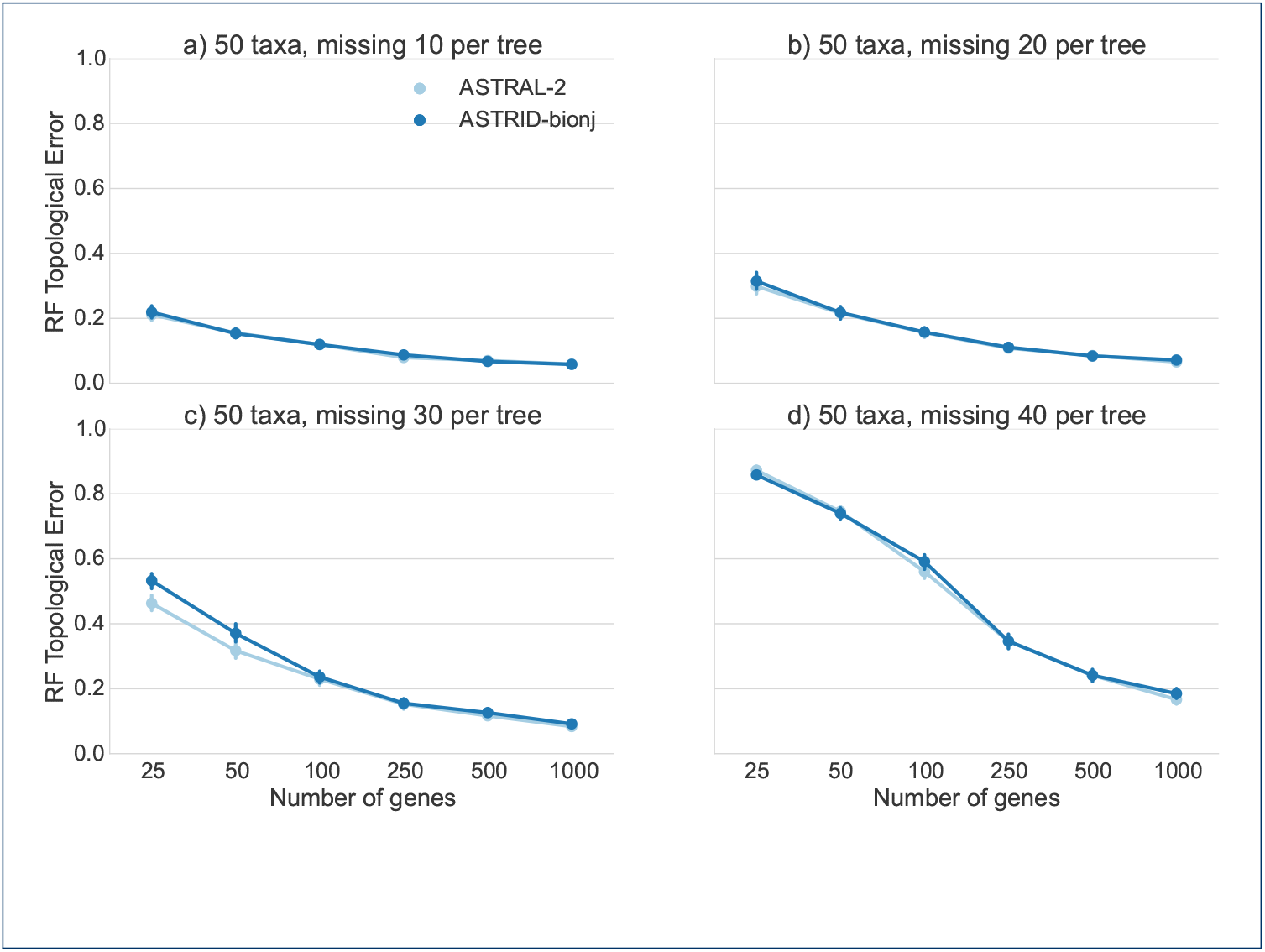
Results on 50-taxon ASTRAL-2 dataset (MC8) with missing taxa and sequence lengths of 150bp. We report RF rates and standard error over 50 replicates.

### Analysis of the mammalian biological dataset

We analyzed the mammalian biological dataset originally studied in [35]. The original dataset had 37 species and 447 genes, but there were 23 erroneous genes (as noted by [20]) which we removed before doing the analysis.

We obtained maximum likelihood gene trees and bootstrap replicates of these gene trees from [22]. We then analyzed these data using ASTRAL-2 and ASTRID+FastME and compared these analyses to previously published trees obtained using ASTRAL and MP-EST [7]. We then annotated the branches of the ASTRID+FastME and ASTRAL-2 trees with bootstrap support from 100 multi-locus bootstrapping (MLBS). The ASTRID+FastME and ASTRAL-2 trees were topologically identical to the ASTRAL tree and differed only in the bootstrap support; see Figure 11 for the ASTRID+FastME tree. On the other hand, the support for the placement of Scandentia - one of the major open questions about mammalian evolution - was very low, only 47% (ASTRAL-2 gave it 82%). Hence, neither the ASTRID tree nor the ASTRAL-2 tree resolved the placement of Scandentia with high support.

**Figure 11.**
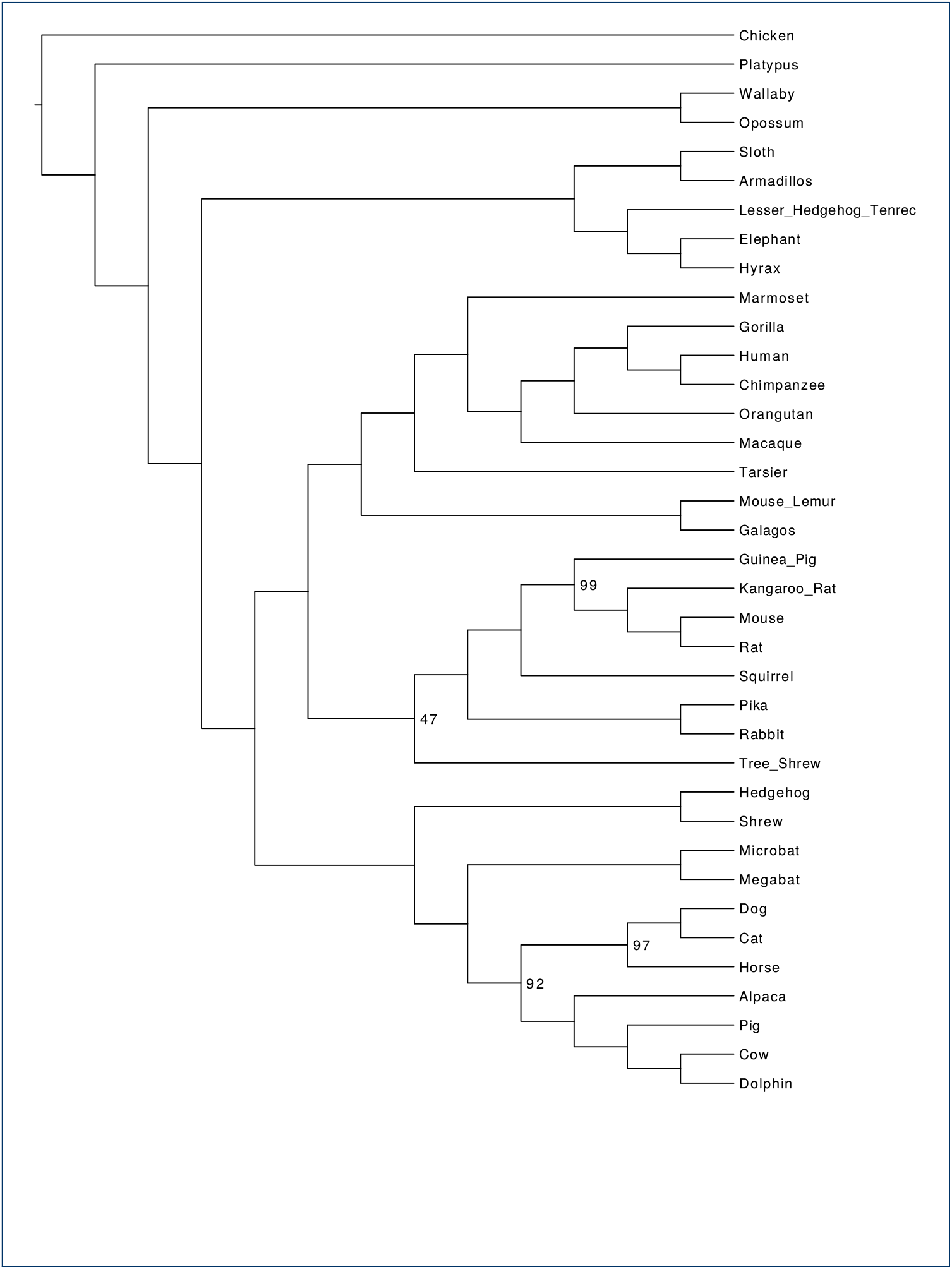
ASTRID analysis of a mammalian biological dataset. We used ASTRID+FastME to analyze the mammalian biological dataset studied in [20, 7], with 37 taxa and 424 genes. The branches are annotated with bootstrap support values from 100 MLBS bootstrap samples; values not shown indicate 100% support. The ASTRID tree is identical to the ASTRAL and ASTRAL-2 trees on the same data, but differs from the MP-EST analysis in the placement of Scandentia.

### Running time results

#### Asymptotic running time

ASTRID has two steps: the first step computes the distance matrix, and the second step uses a selected distance-based method to construct a tree from the distance matrix. When the input has *n* species and *k* genes, then calculating the distance matrix can be performed in *O*(*kn*^2^) time. Distance-based tree estimation methods typically run in *O*(*n*^2^) to *O*(*n*^3^) time, but this step no longer depends on *k*. Hence, the overall running time depends on the selected distance-based method, but is generally dominated by the first phase, especially for typical inputs, for which *k* >> *n*. Thus, under the assumption that *k > n* and that ASTRID uses a distance-based method that runs in *O*(*n*^3^) time, ASTRID’s running time is *O*(*kn*^2^).

ASTRAL-2’s scaling is more complicated to discuss. Asymptotically, ASTRAL-2 runs in *O*(*nk*|*X*|^2^) time, where *n* is the number of species, *k* is the number of genes, and *X* is a set of bipartitions it computes to constrain the search space. The size of *X* is not bounded by a polynomial in the input size, and the technique that ASTRAL-2 uses means that *X* can be large under conditions with high ILS. Thus the asymptotic running times of ASTRAL-2 and ASTRID (used with various distance methods) are quite different.

### Running times on simulated data

In practice, creating the distance matrix took the majority of the running time. On 1000 taxa, creating the distance matrix took several minutes to several hours, depending on the number of genes, but running *F AST M E* took less than one second regardless of the number of genes. However, PhyD* methods were much slower than *F AST M E*; on 1000 taxa, running any of the PhyD* methods took approximately 40 minutes (data not shown).

We recorded running times for ASTRAL-2, ASTRID-FastME, and NJst, on avian simulated datasets with high ILS (1X), as we varied the number of genes (see Fig. 12). Note that ASTRID-FastME was by far the fastest of the three methods, and NJst was the slowest. However, the trends suggest that NJst will be faster than ASTRAL-2 for larger numbers of genes. Note also that ASTRID-FastME and NJst both scaled linearly with the number of genes, but that ASTRAL-2’s running time scaled super-linearly.

**Figure 12.**
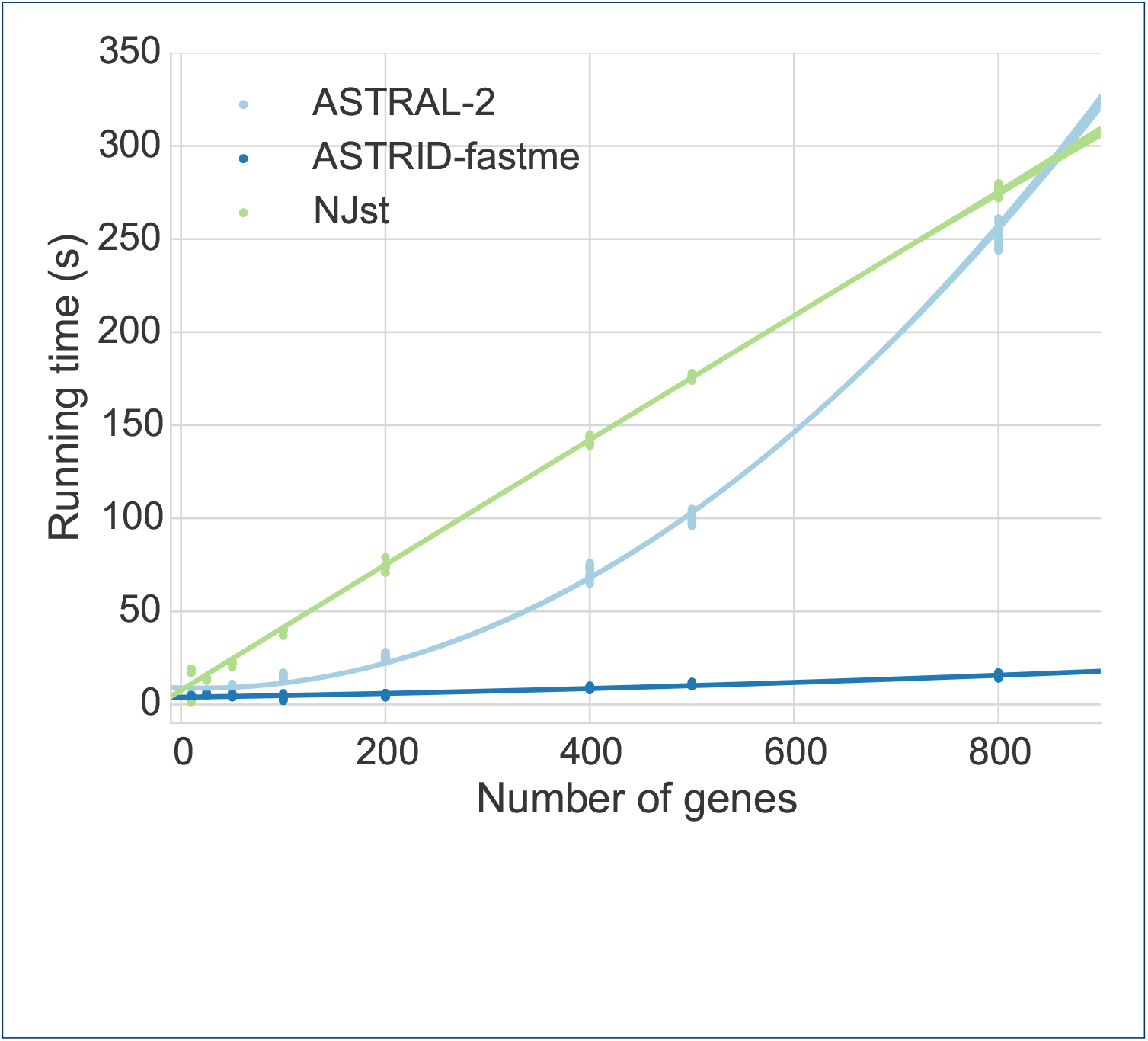
Scatterplot of running times for ASTRID-FastME, ASTRAL-2, and NJst, on avian high ILS (1X) simulated datasets, varying number of genes. We show running time for 20 replicates of each number of genes. The quadratic dependence of ASTRAL-2’s running time is clearly contrasted with the linear dependence of both ASTRID and NJst. Experiments were run on a single core of a 2.7 GHz Intel Xeon processor.

We recorded running times for two variants of ASTRID (one using FastME and the other using BioNJ*), and compared them to ASTRAL-2 on ASTRAL-2 simulated datasets with 1000 taxa (MC11) as we varied the number of genes (Fig. 13) and for 500-gene datasets in which we varied the number of taxa (MC 2 and 7-10, see Fig. 14). The relative running times show that all methods were very fast for smaller datasets, but were clearly distinguished on the larger datasets, where ASTRID-FastME was much faster than ASTRID-BioNJ* and both variants of ASTRID were much faster than ASTRAL-2. For example, on the dataset with 1000 genes and 1000 taxa, ASTRID-FastME finished in 33 minutes, ASTRID-BioNJ finished in 1 hour and 10 minutes, and ASTRAL-2 finished in 12 hours and 30 minutes.

**Figure 13.**
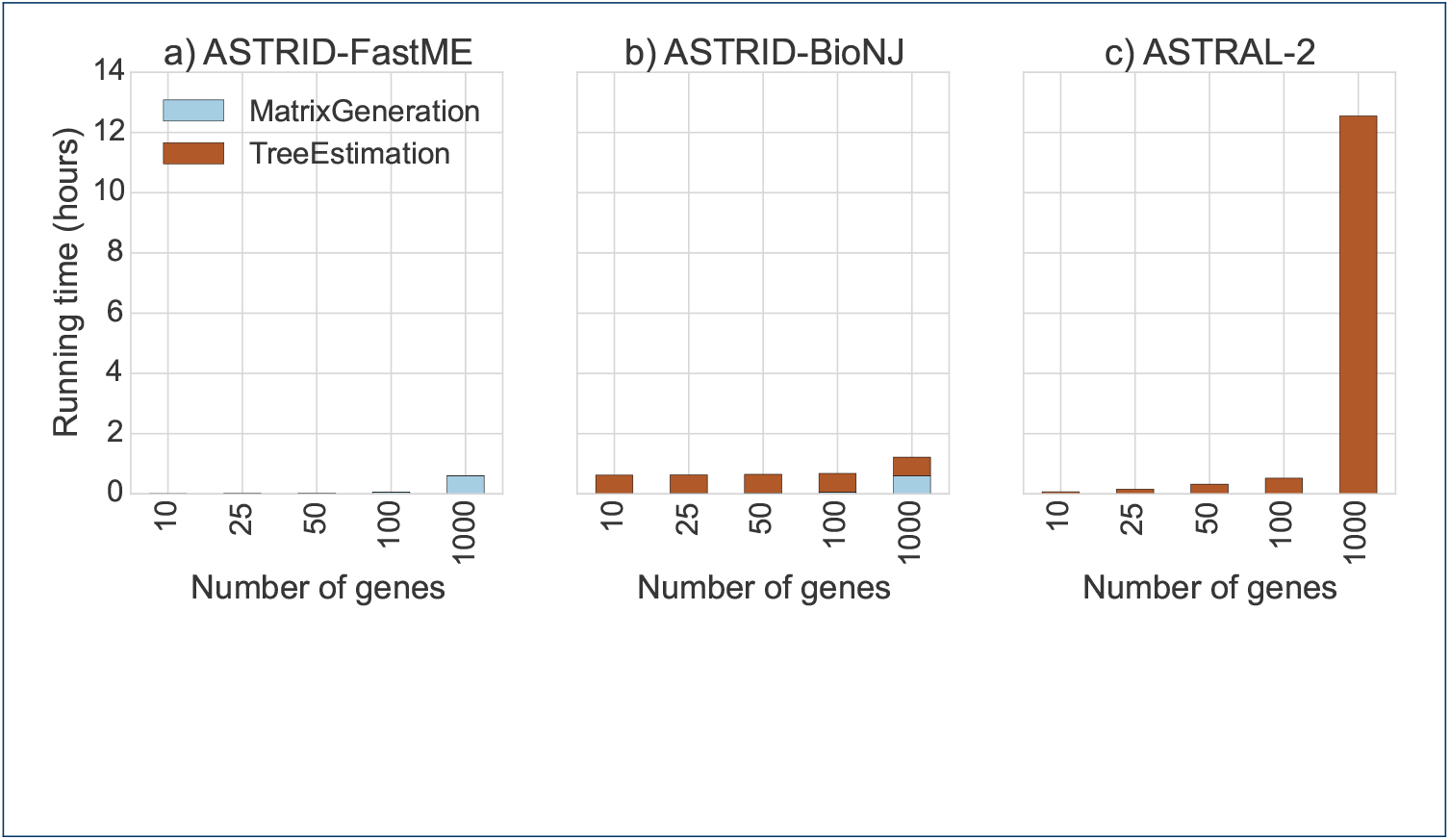
Running time on the ASTRAL-2 simulated datasets with 1000 taxa (MC11), varying number of genes. We show results for the each of the ASTRID steps – matrix generation and tree estimation. We compare ASTRID used with two ways of computing the trees: FastME and BioNJ*. Experiments were run on a single core of a 2.7 GHz Intel Xeon processor.

**Figure 14.**
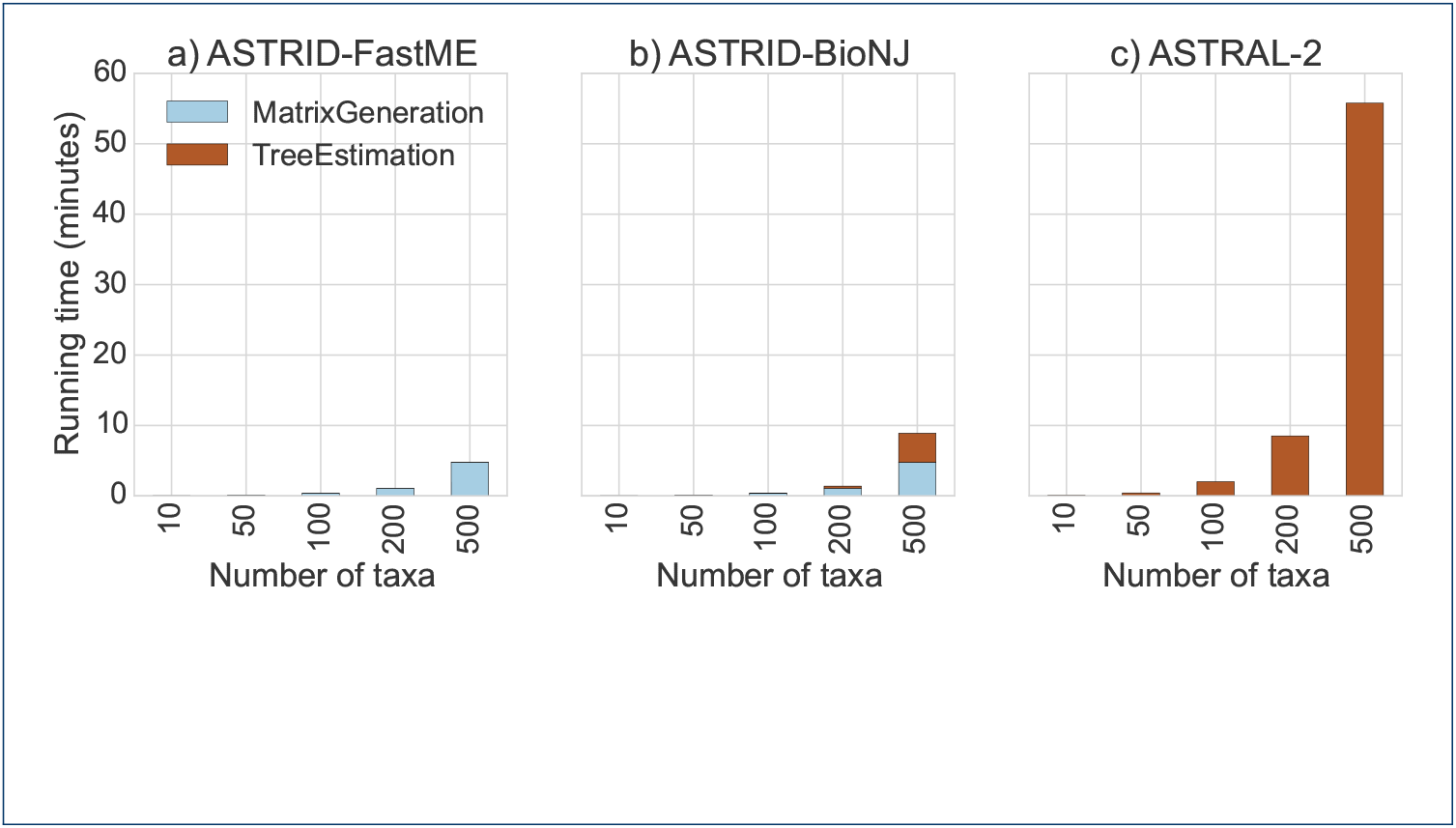
Running time of two ASTRID methods on the ASTRAL-2 simulated dataset with 500 genes, varying number of taxa. We show results on a single replicate of model conditions MC2 and 7-10 from the ASTRAL-2 collection. Experiments were run on a single core of a 2.7 GHz Intel Xeon processor.

### Running times on biological data

We recorded running times for ASTRID-FastME and ASTRAL-2 on the mammalian biological dataset. Both methods took 6 seconds for a single bootstrap replicate on one core of a 2.7 GHz Intel Xeon processor with 424 genes and 37 taxa.

## Discussion

A few trends are apparent upon examining the data as a whole. ASTRAL-2 and ASTRID had, for the most part, very similar levels of accuracy, while MP-EST was consistently less accurate. However, there were cases where ASTRID and ASTRAL-2 have small but detectably different levels of accuracy. One intriguing trend in the data is the improvement of ASTRAL-2 over ASTRID on high ILS datasets; see Figures 6, 7, 8, and 9. In particular, Figures 6 and 7 suggest that increases in ILS should favor ASTRAL-2 over ASTRID. Yet, ASTRID is consistently at least as accurate as ASTRAL-2 on the avian datasets, which have moderate to very high levels of ILS (Fig. 4). Thus, ILS level might have an impact on the relative accuracy of the two methods, but it is not a determining favor. Similarly, neither method dominates the other based on the number of taxa, number of genes, or amount of gene tree estimation error. Thus, it is very difficult to characterize the conditions under which each method is likely to have an advantage over the other. However, even for the cases where there are differences in accuracy, in general the differences are fairly small. Thus, the main difference between the two methods is computational efficiency, where ASTRID is clearly faster. ASTRID has the biggest running time advantage over ASTRAL-2 for large numbers of gene trees, since ASTRID scales linearly in the number of genes while ASTRAL scales super-linearly. This makes ASTRID an especially good method for genome-scale datasets that have a large number of genes.

## Conclusion

ASTRID is a fast and highly accurate method for species tree estimation that is robust to high levels of ILS, and provably statistically consistent under the multispecies coalescent model. Like ASTRAL-2, ASTRID can analyze datasets with unrooted gene trees, an advantage that the two methods have over many other methods (e.g., MP-EST) that can only be run on rooted gene trees. ASTRID (like NJst) runs in time that is polynomial in the number of gene trees and species, but ASTRAL-2 and other leading coalescent-based methods do not have this guarantee. Thus, ASTRID has many desirable theoretical properties compared to existing methods. From an empirical viewpoint, ASTRID is also extremely fast and can analyze very large datasets in minutes, where other methods either cannot run or take hours. In particular, ASTRID is much faster than ASTRAL-2, especially on datasets with many genes and large numbers of species. ASTRID also produces more accurate trees than MP-EST and NJst, and is competitive with ASTRAL-2 in terms of accuracy.

However, even better (more accurate) results might be obtained through more extensive modifications to the ASTRID algorithm design. In particular, the accuracy of the tree depends on the particular distance-based method that is used. New distance-based phylogeny estimation methods, such as the absolute fast converging methods [36, 37, 38, 39], might provide improved accuracy for very large datasets. Another important direction is developing additional methods for estimating species trees from distance matrices that have good accuracy when the distance matrix has missing data. As we saw here, FastME produced more accurate trees than the PhyD* methods, but it could only be applied to distance matrices without any missing data. An extension of FastME to enable it to handle incomplete distance matrices would also be of great interest.

This study can be expanded in several directions. Future work should more carefully investigate the conditions under which ASTRID is more reliable than ASTRAL-2, and explore performance on more biological datasets. This study also only investigated relatively long sequences; a subsequent study should investigate the relative and absolute accuracy of ASTRID and other methods on very short sequences, since recombination-free loci can be very short [32]. In addition, this study only examined datasets with a single individual per species, yet ASTRID (like NJst) can be run on datasets with multiple individuals; future work should evaluate the absolute and relative accuracy of ASTRID and other methods on such data. This study showed that ASTRID performed well in terms of species tree topology estimation, but we did not explore its accuracy with respect to the estimation of coalescent branch lengths; future work will need to explore how well ASTRID estimates these numeric parameters. Finally, it may well be that ASTRID will be most useful as a starting tree for use within more computationally intensive analyses, including Bayesian MCMC analyses (e.g., *BEAST) or maximum likelihood analyses.

## Availability of supporting data

All datasets used in this study are available from prior publications. ASTRID is available in open source form on github at http://pranjalv123.github.io/ASTRID. Supporting materials are available online at http://pranj.al/ASTRID.

## Abbreviations

AD: Average distance
ILS: Incomplete lineage sorting
MSC: Multi-species coalescent
RF: Robinson-Foulds

## Author’s contributions

PV implemented ASTRID, performed experiments, wrote the first draft, and analyzed the data. TW designed the study, analyzed the data, and wrote the final draft.

## Acknowledgements

PV was supported by the Roy J. Carver graduate fellowship from the University of Illinois at Urbana-Champaign, College of Engineering. TW was supported by National Science Foundation grant DBI-1461364. PV and TW were both supported by the College of Engineering at the University of Illinois at Urbana-Champaign through a gift from the Grainger Foundation.

